# Extracellular Degradation into Adenosine and the Activities of Adenosine Kinase and AMPK Mediate Extracellular NAD^+^-produced increases in the Adenylate Pool of BV2 Microglia under Basal Conditions

**DOI:** 10.1101/334268

**Authors:** Jie Zhang, Caixia Wang, Danhong Wu, Weihai Ying

**Affiliations:** Med-X Research Institute and School of Biomedical Engineering, Shanghai Jiao Tong University, Shanghai 200030, P.R. China; Department of Otorhinolaryngology, Shanghai Sixth People’s Hospital, Shanghai Jiao Tong University, Shanghai 200233, P.R. China; Department of Neurology, Shanghai Fifth People’s Hospital, Fudan University, Shanghai, P.R. China

**Keywords:** NAD+, Microglia, Adenylate Pool, Adenosine, AMPK

## Abstract

Cumulating evidence has indicated NAD^+^ deficiency as a common central pathological factor of multiple diseases and aging. NAD^+^ supplement is highly protective in various disease and aging models, while two key questions remain unanswered: 1) Does extracellular NAD^+^ also produce its effects through its degradation product adenosine? 2) Does extracellular NAD^+^ produce the protective effects by affecting cells under pathological insults only, or by affecting both normal cell and cells under pathological insults? Since extracellular NAD^+^ can be degraded into adenosine, and endogenous adenosine levels are in the nanomolar range under physiological conditions, extracellular NAD^+^ may produce its effects through its degradation into adenosine. In this study we used BV2 microglia as a cellular model to test our hypothesis that NAD^+^ treatment can increase the intracellular adenylate pool under basal conditions through its extracellular degradation into adenosine. Our study has shown that extracellular NAD^+^ increases the adenylate pool of BV2 microglia under basal conditions through its degradation into adenosine that enters the cells through equilibrative nucleoside transporters. The intracellular adenosine is converted to AMP by adenosine kinase, which increases intracellular ATP by both activating AMPK and increasing ADP that drives mitochondrial FoF1-ATP synthase. Collectively, our study has suggested that extracellular NAD^+^ can enhance defensive capacity of normal cells through a novel pathway, which includes extracellular NAD^+^ degradation into adenosine and the activities of adenosine kinase and AMPK. Our findings have also suggested that NAD^+^ administration in various disease and aging models may significantly affect the microglia under basal conditions.

## Introduction

NAD^+^ plays critical roles in a number of biological functions including energy metabolism, mitochondrial activity, DNA repair and cell death (Ying, 2008). Accumulating evidence has also suggested that there are significant decreases in the NAD^+^ levels in the models of aging and a number of diseases (Ying, 2008; Verdin, 2015; Zhang and Ying, 2018). Supplement with NAD^+^ can produce profound beneficial effects in animal models of aging and multiple diseases (Ying et al., 2007; Yoshino et al., 2011; Mills et al., 2016; Zhang et al., 2016; Das et al., 2018). Collectively, these findings have indicated that NAD^+^ deficiency is a common central pathological factor of a number of diseases and aging (Ying, 2008; Imai and Guarente, 2014; Verdin, 2015; Zhang and Ying, 2018).

While NAD^+^ supplement can produce beneficial effects in various models of diseases and aging (Weihai Ying, 2007; Yoshino et al., 2011; Mills et al., 2016; Zhang et al., 2016; Das et al., 2018), two key questions regarding the mechanisms underlying the protective effects remains unanswered: 1) Does extracellular NAD^+^ produce its effects through its degradation products such as adenosine and nicotinamide? 2) Does extracellular NAD^+^ produce its protective effects by affecting cells under pathological insults only, or by affecting both normal cells and cells under pathological insults?

Some previous findings have provided answers to the first question: Intranasal administration of NAD^+^, but not nicotinamide, can profoundly decrease ischemic brain injury (Ying et al., 2007) and traumatic brain injury (Won et al., 2012), which argues against the possibility that extracellular NAD^+^ produces its protective effects via nicotinamide – one of its degradation products. However, extracellular NAD^+^ can also be degraded into adenosine by multiple potential mechanisms (Moreschi et al., 2006; Klein et al., 2009; Okuda et al., 2010b; Adriouch et al., 2012). While it has been reported that the radiolabeled adenosine moiety was observed in astrocytes within 10 mins of treatment of the cells with radiolabeled NAD^+^ (Okuda et al., 2010a), there has been no report that provides answers to the following question: Does extracellular NAD^+^ produce its effects through adenosine? Since it is established that adenosine can produce beneficial effects under multiple pathological conditions (Ely and Berne, 1992; Zhang et al., 1993; Yap and Lee, 2012), in our current study we used BV2 microglia as a cellular model to test our hypothesis that extracellular NAD^+^ produces its biological effects on cells via its extracellular degradation into adenosine.

Regarding the second key question, there are studies suggesting that extracellular NAD^+^ produces its protective effects at least partially by directly entering cells to decrease the damage of the cells under pathological insults: First, because such insults as oxidative stress can lead to rapid decreases in the cytosolic NAD^+^ concentrations by activating PARP-1 (Ying et al., 2003; Alano et al., 2004; Pillai et al., 2005; Alano et al., 2010), it is reasonable to propose that extracellular NAD^+^ may enter cells via P2X7 receptors (Ying et al., 2003; Alano et al., 2004; Pillai et al., 2005; Alano et al., 2010) or Connexin 43 hemichannels (Bruzzone et al., 2001) by gradient-dependent mechanisms; second, NAD^+^ administration can significantly enhance the NAD^+^ levels of the tissues under various pathological insults (Ying, 2008; Imai and Guarente, 2014; Verdin, 2015; Zhang and Ying, 2018); and third, numerous studies have reported that NAD^+^ treatment can decrease the death of various types of cells exposed to oxidative stress, DNA alkylating agents or excitotoxins (Ying et al., 2003; Alano et al., 2004; Pillai et al., 2005; Alano et al., 2010). The major mechanisms underlying these beneficial effects include improvement of glycolysis by restoring the activity of the NAD^+^-dependent enzyme glyceraldehyde-3-phosphate dehydrogenase (GAPDH), prevention of mitochondrial depolarization and mitochondrial permeability (MPT), activation of SIRT1 and SIRT3, and promotion of DNA repair (Ying et al., 2003; Alano et al., 2004; Pillai et al., 2005; Ying, 2008; Alano et al., 2010; Zhang and Ying, 2018).

However, there has been little information suggesting that the NAD^+^ administration may also decrease tissue damage by enhancing the defensive potential of normal cells that have not been attacked at the time of exposures to the administered NAD^+^. Since the intracellular NAD^+^ concentrations in normal cells range from 1 mM – 10 mM (Ying, 2008), it is unlikely that the NAD^+^ administration, usually at the doses between 10 mg – 20 mg/kg, may produce NAD^+^ concentrations in the blood which surpasses 1 mM. Therefore, it is unlikely that extracellular NAD^+^ may directly enter normal cells to produce its effects in these models. However, since endogenous adenosine levels are in the nanomolar range under physiological conditions (Latini and Pedata, 2001), it is possible that the adenosine that is generated from extracellular NAD^+^ degradation may enter cells under basal conditions through equilibrative nucleoside transporters (ENTs) to produce various biological effects.

There have been few studies regarding the effects of NAD^+^ treatment on cellular properties under basal conditions. Our recent study has shown that NAD^+^ treatment can induce significant increases in the intracellular ATP levels of BV2 microglia under basal conditions, while the mechanisms underlying this effect of NAD^+^ require future studies (Zhang et al., 2018). Because intracellular ADP and AMP are closely related with intracellular ATP (Stryer, 1995), we propose that NAD^+^ treatment can significantly increase the intracellular adenylate pool of BV2 microglia under basal conditions.

Increasing evidence has indicated that inflammation plays critical pathological roles in multiple neurodegenerative disorders (Frank-Cannon et al., 2009; Hirsch and Hunot, 2009). Microglial activation is a key event of neuroinflammation in neurodegenerative diseases, which can generate oxidative stress and cytokines thus impairing neuronal survival (Frank-Cannon et al., 2009; Hirsch and Hunot, 2009). Elucidation of the mechanisms underlying the regulation of functions and survival of both activated microglia and microglia under resting conditions is of critical significance for understanding the roles of microglia in neurodegenerative disorders and brain aging. Previous studies have indicated that ATP plays critical roles in both microglia under resting conditions and activated microglia: Basal ATP release is necessary for its survival in microglia; and extracellular ATP induces migration, chemotaxis and ramification of microglia (Honda et al., 2001; Wollmer et al., 2001; Duan et al., 2009; Ma et al., 2014). Therefore, studies on the effects of NAD^+^ treatment on intracellular levels of adenylate of microglia can significantly enhance our understanding on the mechanisms underlying the effects of NAD^+^ treatment in various models of neurological diseases and aging.

In this study we used BV2 microglia as a cellular model to test our hypothesis that extracellular NAD^+^ degradation into adenosine mediates the effects of NAD^+^ treatment on cells under basal conditions. Our study has provided multiple lines of evidence suggesting that extracellular NAD^+^ produces increased adenylate pool of BV2 microglia under basal conditions by the mechanisms including such key factors as extracellular degradation into adenosine and activities of adenosine kinase and AMPK.

## Materials and methods

### Reagents

NADH (N4505), NAD^+^ (N0632), dorsomorhpin (P5499), Adenosine 5′-monophosphate (AMP, 01930), pyridoxyl phosphate-6-azophenyl-2′,4′-disulfonic acid (PPADS, P178), 5-Iodotubercidin (I-100) and dipyridamole (DPR, D9766) were purchased from Sigma Aldrich (St. Louis, MO, USA). AMPK siRNAs and control siRNA were purchased from GenePharma (Shanghai, China).

### Cell culture

BV2 microglia was obtained from Institute of Neurology, Ruijin Hospital (Shanghai, China). The cells were plated into 24-well culture plates at the initial density of 6×10^5^ cells/mL in Dulbecco’s Modified Eagle Medium (HyClone, Logan, Utah, USA) containing 5% fetal bovine serum (Gibco, Carlsbad, CA), 100 units/ml penicillin and 100 μg/ml streptomycin. The cells were maintained at 37°C in 5% CO_2_ incubator.

### RNA interference

BV2 microglia were transfected with either AMPK siRNA oligonucleotides (5‘ GAGAAGCAGAAGCACGACGTT 3‘) or control siRNA oligonucleotides (5‘ UUCUCCGAACGUGUCACGUTT 3‘) (Genepharma, Shanghai, China), when the cells were approximately 50% confluent. Lipofectamine 2000 (Invitrogen, Carlsbad, CA, USA) was used for the transfection according to the manufacturer’s instructions. For each well of a 24-well plate, 100 µl Opti-MEM containing 0.06 nmol of the siRNA oligonucleotides and 2 µl lipofectamine 2000 was added into 500 µl culture media of the cells. The AMPK protein level was determined by Western Blot 18 h after the transfection.

### Western Blot assays

BV2 microglia was washed by cold PBS and lysed in RIPA buffer (Millipore, Temecula, CA, USA) containing Complete Protease Inhibitor Cocktail (Roche Diagnostics, Mannheim, Germany) and 1 mM PMSF. BCA Protein Assay Kit (Pierce Biotechonology, Rockford, IL, USA) was used to determine the concentrations of the protein samples. Thirty μg of total protein was electrophoresed through a 10% SDS-PAGE gel, and then transferred to 0.45 μm nitrocellulose membranes. The membranes were incubated with 5% (v/v) BSA for 2 h. The blots were incubated overnight at 4 °C with rabbit-derived anti-AMPKα antibody (Cell Signaling Technology, Danvers, MA, USA) or goat-derived anti-actin antibody (Santa Cruz Biotechnology, CA, USA), then incubated with HRP-conjugated secondary antibody (Epitomics, Hangzhou, Zhejiang Province, China). Protein signals were visualized using the ECL detection reagent (Pierce Biotechnology, Rockford, IL, USA). The intensities of the bands were analyzed by densitometry using computer-based Image-Pro Plus program.

### Assays of ATP, ADP and AMP

Adenine nucleotides were determined by a luciferin-luciferase assay adapted from Spielmann et al. (Spielmann et al., 1981). In brief, cells were harvested with 200 μL Tris-EDTA buffer (0.1 M Tris–HCl, pH 7.75 and 1 mM EDTA) and heated at 95 °C for 5 minutes. The ATP contents were determined using the luciferin and luciferase from an ATP Bioluminescence Assay Kit (Roche Applied Science, Mannheim, Germany). ADP and AMP were also quantified after their conversion into ATP by pyruvate kinase and myokinase, respectively. The amount of ADP was obtained by subtracting the ATP value from the (ATP^+^ADP) value, and the amount of AMP was calculated from the difference between the (ATP^+^ADP) content and the (ATP^+^AMP^+^ADP) content.

### Adenosine Level Assay

Cells were collected, resuspended in PBS buffer and went through three freeze-thaw cycles. After centrifugation at 3000 RPM for 20 min at 4 □, the supernatant was collected for use. The adenosine level was measured by adenosine assay kit (Biovision, CA, USA) according to the manufacturer’s protocol. The fluorescent intensity was detected at Ex/Em = 535/587 nm.

### NAD^+^ assay

NAD^+^ concentrations were measured by recycling assay as described previously (Ying et al., 2003). Briefly, cells were washed with cold PBS for once and extracted in 0.5 N cold perchloric acid. After centrifugation at 12,000 RPM for 5 min at 4 °C, the supernatant was neutralized to pH 7.2 using 3 N KOH and 1 M potassium phosphate buffer. After centrifugation at 12,000 RPM for 5 min at 4 °C, 50 μl supernatant was mixed with 100 μl reaction medium containing 0.0425 mg/ml 3-[4,5-dimethylthiazol-2-yl]-2,5-diphenyl-tetrazolium bromide (MTT), 0.0296 mg/ml phenazine methosulfate, 0.0325 mg/ml alcohol dehydrogenase, 12.21 mg/ml nicotinamide, and 6% (v/v) ethanol in Gly-Gly buffer (65 mM, pH 7.4). After 10 min, the optical absorbance of the samples at 556 nm was measured with a plate reader.

### Mitochondrial membrane potential assay

Mitochondrial membrane potential (Δψ_m_) was determined by flow cytometry-based JC-1 (5,5’,6,6’-tetrachloro-1,1’,3,3’-tetraethyl-benzimidazolylcarbocyanine iodide) assay according to the manufacturer’s instruction. Cells were harvested by 0.25% trypsin-EDTA, and incubated in cell media containing 5 μg/mL JC-1 (Enzo Life Sciences, NY, USA) for 15 min at 37 °C. After washed once with PBS, samples was analyzed by a flow cytometer (FACSAria II, BD Biosciences, NJ, USA) using the excitation wavelength of 488 nm and the emission wavelengths of 525 nm for green fluorescence, or the emission wavelengths of 575 nm for orange-red fluorescence. The Δψ_m_ of each cell was calculated by the ratio of red fluorescence intensity to green fluorescence intensity. For each sample, the ratio of the cells with low Δψ_m_ in total cell population was reported by the flow cytometer (FACSAria II, BD Biosciences, NJ, USA).

### Glycolytic rate assay

2-NBDG (2-(N-(7-Nitrobenz-2-oxa-1,3-diazol-4-yl)Amino)-2-Deoxyglucose) is a fluorescent glucose analog that has been used to monitor glucose uptake in live cells. BV2 microglia were incubated in media containing 100 μM 2-NBDG (Invitrogen, Carlsbad, CA, USA) for 1 hrs in 37 °C, then harvested by 0.25% trypsin-EDTA. After washed once with PBS, BV2 microglia was analyzed by a flow cytometer r (FACSAria II, BD Biosciences, NJ, USA), which detected emission fluorescence at the wavelength of 518 nm (FITC-A) with excitation wavelength of 488 nm.

### Statistical analysis

Data were presented as mean ± SEM and analyzed by ANOVA followed by Student-Newman-Keuls *post hoc* test. *P* values less than 0.05 were considered statistically significant.

## Results

### NAD^+^ treatment can increase the intracellular levels of ATP, ADP, AMP and NAD^+^ of BV2 microglia under basal conditions

We determined the effects of NAD^+^ on the intracellular adenylate levels of BV2 microglia, showing that treatment of 10, 100 and 500 μM NAD^+^ can significantly increase the intracellular levels of ATP, ADP and AMP of the cells (Figures 1A, 1B, 1C). The NAD^+^ treatment also significantly increased the intracellular NAD^+^ levels of the cells (Figure 1D).

**Figure 1:**
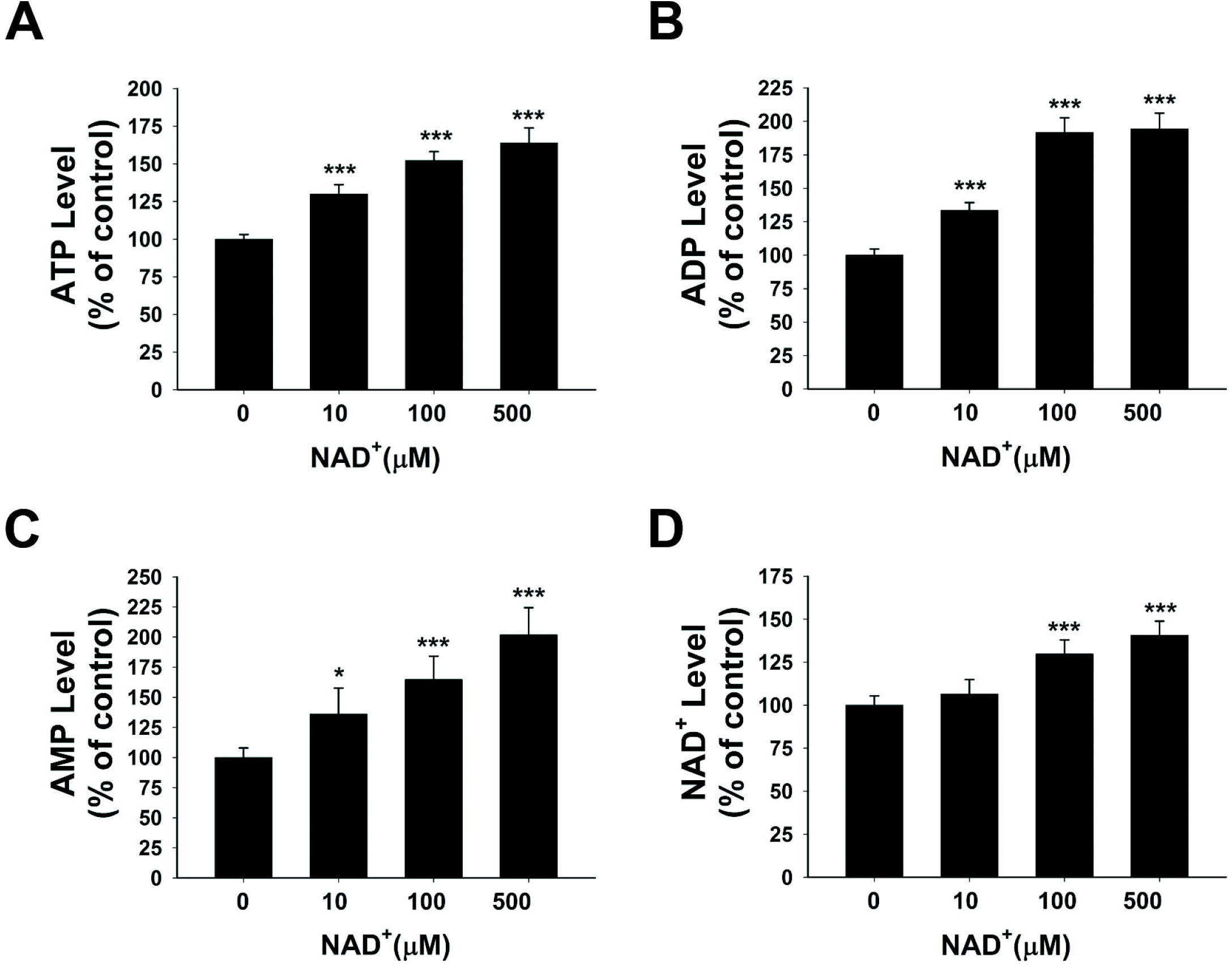
Effects of NAD^+^ treatment on the intracellular levels of ATP, ADP, AMP and NAD^+^ of BV2 microglia under basal conditions. (A) NAD^+^ treatment significantly increased the intracellular ATP level of the cells. (B) NAD^+^ treatment significantly increased the intracellular ADP level of the cells. (C) NAD^+^ treatment significantly increased the intracellular AMP level of the cells. (D) NAD^+^ treatment significantly increased the intracellular NAD^+^ level of the cells. BV2 cells were treated with NAD^+^ for 3 h before the assays were conducted. N = 16. The data were pooled from four independent experiments. *, *P* < 0.05; **, *P* < 0.01; ***, *P* < 0.001.

### Roles of glycolytic rate, mitochondrial membrane potential and SIRT1 in the NAD^+^ treatment-induced increases in the adenylate pool of BV2 microglia under basal conditions

Glycolysis and mitochondrial oxidative phosphorylation (OXPHOS) are major pathways for ATP production, in which NAD^+^ plays significant roles (Stryer, 1995). Previous studies have suggested that NAD^+^ treatment decreases cell death induced by oxidative stress, alkylating agents and excitotoxins by such mechanisms as improving glycolysis, preventing mitochondrial depolarization and activating SIRT1 (Ying et al., 2003; Alano et al., 2004; Pillai et al., 2005; Ying, 2008; Alano et al., 2010; Zhang and Ying, 2018). In order to determine if improving glycolysis and preventing mitochondrial depolarization are also major mechanisms underlying the NAD^+^ treatment-produced increases in the adenylate pool of BV2 microglia under basal conditions, we determined the effects of NAD^+^ treatment on the glycolytic rate and mitochondrial membrane potential of the cells under basal conditions: NAD^+^ treatment did not significantly affect the glycolytic rate of the cells under basal conditions (Supplementary Figure 1A). We also found that NAD^+^ treatment was incapable of affecting the mitochondrial membrane potential of the cells (Supplementary Figures 1B, 1C). Moreover, neither the Complex I inhibitor rotenone nor the Complex IV inhibitor sodium azide was capable of preventing the NAD^+^-induced increases in the intracellular ATP levels (Supplementary Figures 1D, 1E).

We further found that SIRT1 inhibitor EX527 was incapable of affecting the NAD^+^-induced increases in the intracellular ATP levels of the cells (Supplementary Figure 2), thus arguing against the possibility that SIRT1 mediates the NAD^+^ treatment-induced increases in the adenylate pool in our study.

**Figure 2:**
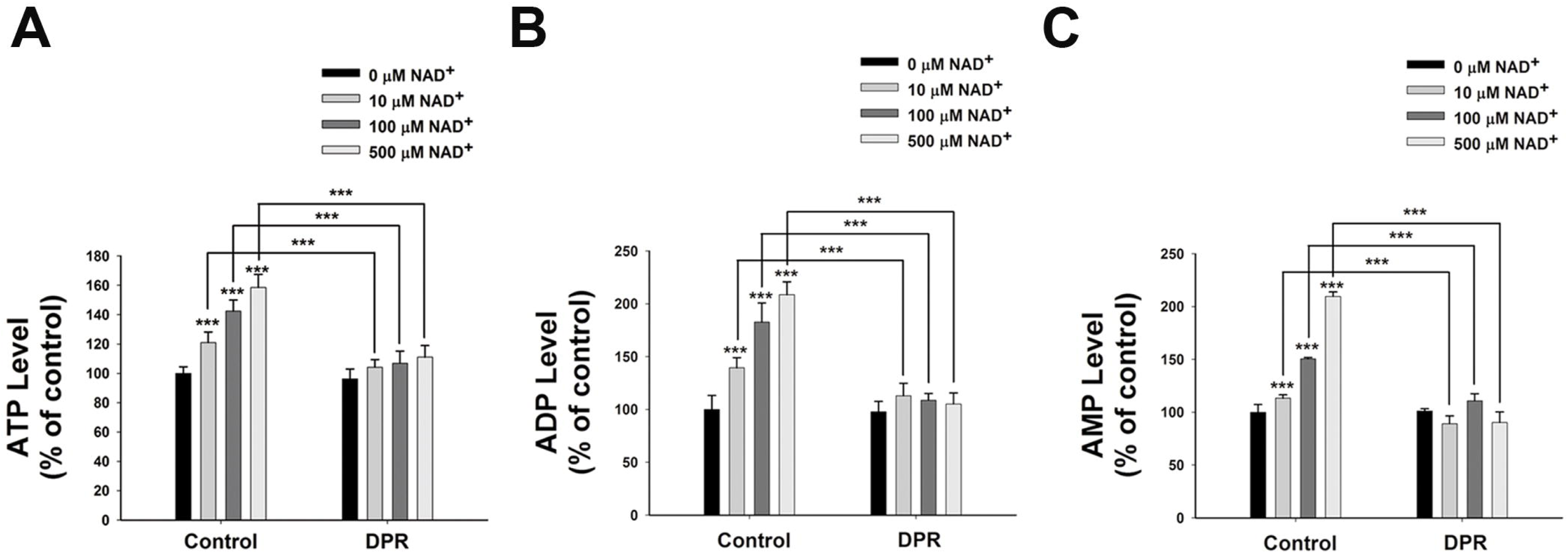
Adenosine uptake plays a key role in the NAD^+^-induced increases in the intracellular levels of ATP, ADP and AMP of BV2 microglia under basal conditions. Treatment of the cells with 0.5 μM DPR completely blocked the NAD^+^-induced increases in the intracellular ATP levels of the cells. (B) Treatment of the cells with 0.5 μM DPR completely blocked the NAD^+^-induced increases in the intracellular ADP levels of the cells. (C) Treatment of the cells with 0.5 μM DPR completely blocked the NAD^+^-induced increases in the intracellular AMP levels of the cells. The cells were co-treated with NAD^+^ and DPR for 3 h. Subsequently the assays on intracellular ATP, ADP and AMP levels were conducted. N=16. The data were pooled from four independent experiments. *, *P* < 0.05; **, *P* < 0.01; ***, *P* < 0.001.

### Contribution of adenosine transport to the NAD^+^-induced increases in the intracellular adenylate levels of BV2 microglia under basal conditions

It has been reported that extracellular NAD^+^ can be degraded into adenosine and nicotinamide (Okuda et al., 2010a). Adenosine can be transported by ENTs into cells (Latini and Pedata, 2001), which may be used for synthesis of AMP, ADP and ATP (Rapaport and Zamecnik, 1978). Due to the relatively high activity of intracellular adenosine kinase and normally low intracellular adenosine levels, the net flux through ENTs is inwardly directed under normal conditions (Latini and Pedata, 2001). To test our hypothesis that extracellular NAD^+^ may lead to increased intracellular adenylate pool by its extracellular degradation into adenosine that is subsequently transported into cells, we determined the effects of dipyridamole (DPR), an inhibitor of ENTs (Wang et al., 2013), on the NAD^+^-induced increases in the intracellular ATP levels. We found that DPR blocked the NAD^+^-induced increases in the intracellular levels of ATP, ADP and AMP (Figures 2A, 2B, 2C).

To exclude the possibility that NAD^+^ produced the effects on the adenylate pool by producing nicotinamide – another degradation product of NAD^+^, we determined if nicotinamide may affect the intracellular ATP levels. Neither 10 nor 100 μM nicotinamide affected the intracellular ATP levels, while 500 μM nicotinamide increased the intracellular ATP levels by merely 10% (Supplementary Figure 3). These observations argued against the possibility that the nicotinamide moiety is responsible for the NAD^+^-induced increases in the adenylate levels.

**Figure 3:**
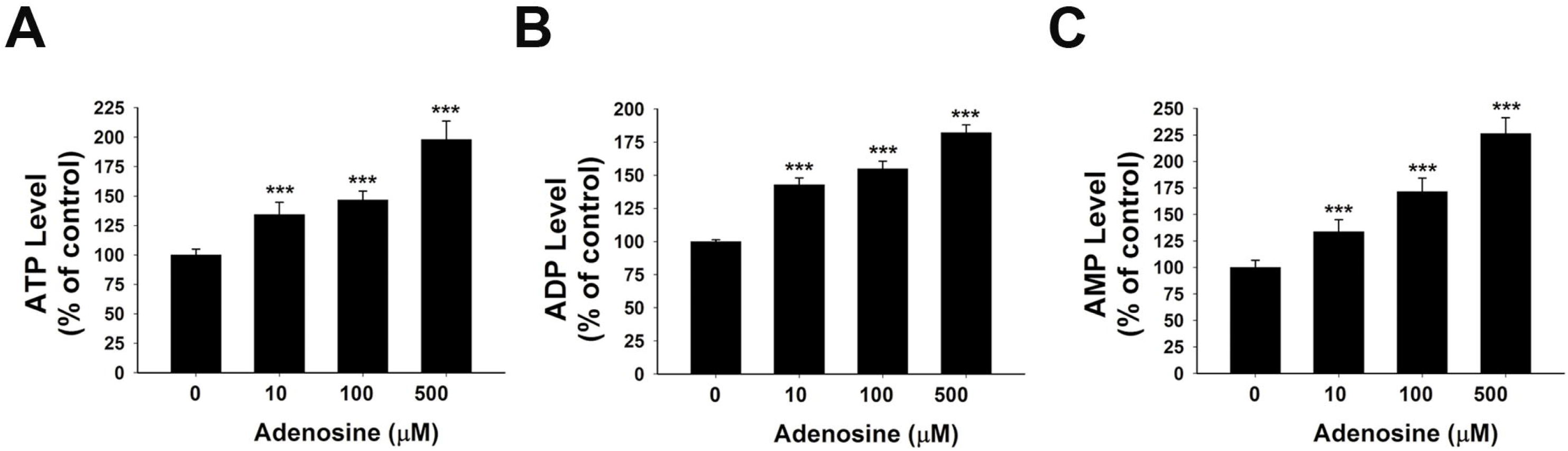
Adenosine treatment led to significant increases in the intracellular levels of ATP, ADP and AMP of BV2 microglia under basal conditions. (A) Treatment of the cells with adenosine significantly increased the intracellular ATP levels of the cells. (B) Treatment of the cells with adenosine significantly increased the intracellular ADP levels of the cells. (C) Treatment of the cells with adenosine significantly increased the intracellular AMP levels of the cells. N=16. The data were pooled from four independent experiments. *, *P* < 0.05; **, *P* < 0.01; ***, *P* < 0.001.

### Extracellular adenosine is capable of increasing the intracellular adenylate pool of BV2 microglia under basal conditions

To further test our proposal that the extracellular adenosine that is generated from extracellular NAD^+^ degradation can produce increased adenylate levels of BV2 microglia under basal conditions, we determined the effects of 10, 100 and 500 μM adenosine treatment on the intracellular ATP, ADP and AMP levels of BV2 microglia under basal conditions. Our study showed that the adenosine at all of these concentrations was capable of significantly increasing the intracellular adenylate levels of the cells (Figures 3A, 3B, 3C).

### Roles of adenosine kinase in the NAD^+^-induced increases in the adenylate levels of BV2 microglia under basal conditions

Because endogenous adenosine levels are in the nanomolar range under normal physiological conditions (Latini and Pedata, 2001), it is possible that the concentrations of the adenosine generated from the degradation of 10 – 500 μM NAD^+^ in our study may surpass the intracellular adenosine concentrations, thus leading to entrance of the extracellular adenosine into the cells. We found that treatment of BV2 microglia with 10, 100 and 500 μM NAD^+^ significantly increased the intracellular adenosine levels (Figure 4A).

**Figure 4:**
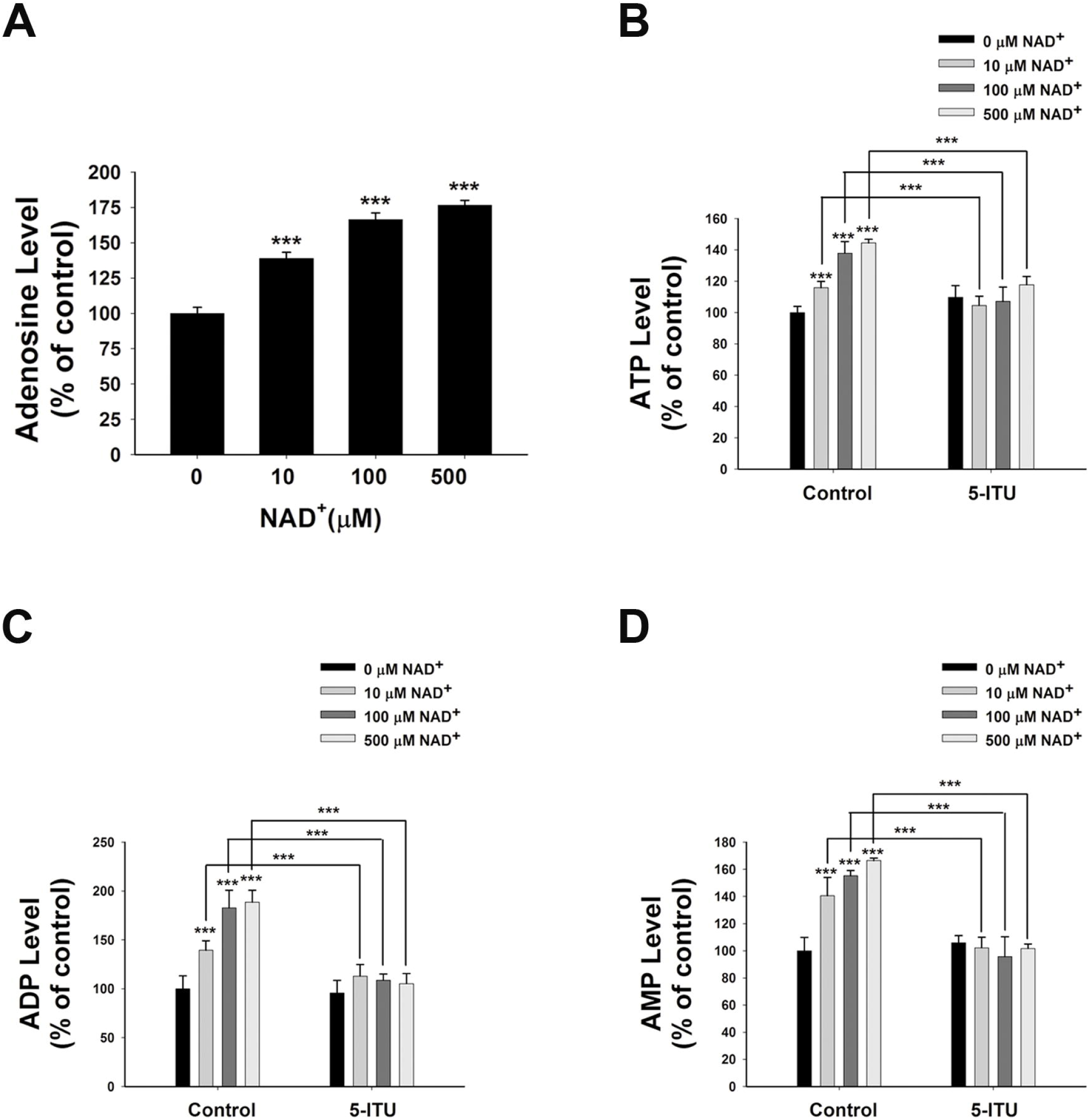
Adenosine kinase mediates the NAD^+^-induced increases in the intracellular levels of ATP, ADP and AMP of BV2 microglia under basal conditions. (A) NAD^+^ treatment significant increased the intracellular adenosine level of the cells. (B) Treatment of the cells with5 μM 5-ITU blocked the NAD^+^-induced increases in the intracellular ATP levels of the cells. (C) Treatment of the cells with 5 μM 5-ITU blocked the NAD^+^-induced increases in the intracellular ADP levels of the cells. (D) Treatment of the cells with 5 μM 5-ITU blocked the NAD^+^-induced increases in the intracellular AMP levels of the cells. The cells were co-treated with 5 μM 5-ITU and NAD^+^ for 3 h before the assays were conducted. N = 16. The data were pooled from four independent experiments. ***, *P* < 0.001.

It has been reported that adenosine can be converted to AMP, which is mediated by adenosine kinase (Miller et al., 1979), leading to AMPK activation (da Silva et al., 2006). Therefore, we applied the adenosine kinase inhibitor 5-Iodotubercidin (5-ITU) to test our hypothesis that the extracellular NAD^+^ can produce increased intracellular AMP, ADP and ATP by increasing intracellular adenosine that is converted to AMP by adenosine kinase. We found that treatment of the cells with 5 μM 5-ITU virtually completely blocked the NAD^+^-induced increases in the intracellular AMP, ADP and ATP levels of BV2 microglia under basal conditions (Figures 4B, 4C, 4D).

### Roles of AMPK in NAD^+^-induced increases in the adenylate levels of BV2 microglia under basal conditions

Since AMPK is a crucial regulator of energy metabolism, which can be regulated by such factors as AMP / ATP ratios (Hardie, 2015), we determined if AMPK is involved in the NAD^+^-induced increases in the intracellular ATP levels of BV2 cells by assessing the effects of NAD^+^ treatment on AMPK phosphorylation. Both 100 and 500 μM NAD^+^ led to increased AMPK phosphorylation (Figure 5A). The AMPK phosphorylation induced by NAD^+^ was blocked by the adenosine kinase inhibitor 5-ITU (Figure 5B). The NAD^+^treatment did not affect the AMP / ATP ratios (Figure 5C), thus arguing against the possibility that the NAD^+^ treatment led to increased AMPK activity by increasing AMP / ATP ratios.

**Figure 5:**
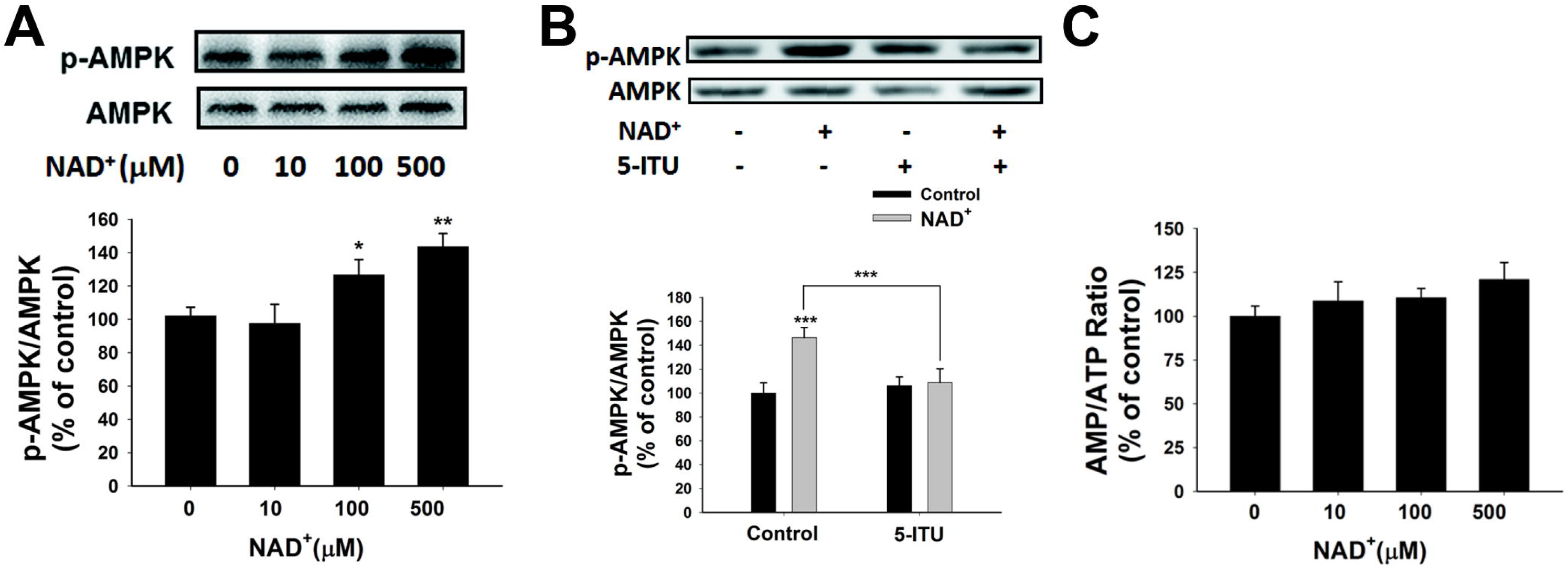
NAD^+^ treatment increased AMPK phosphorylation of BV2 microglia under basal conditions. (A) Both 100 and 500 μM NAD^+^ increased AMPK phosphorylation. Five μM 5-ITU blocked the NAD^+^-induced AMPK phosphorylation in BV2 microglia. NAD^+^ treatment did not affect the AMP / ATP ratios of the cells. N = 12. The data were pooled from four independent experiments. *, *P* < 0.05; **, *P* < 0.01; ***, *P* < 0.001.

We further determined the roles of AMPK in the NAD^+^-induced increases in the intracellular ATP levels, showing that the AMPK inhibitor dorsomorphin blocked the NAD^+^-induced increases in the ATP levels (Figure 6A). Moreover, treatment of the cells with AMPK siRNA, which led to a significant decrease in the AMPK levels of the cells (Figure 6B), also produced significant attenuation of the NAD^+^-induced increases in the intracellular ATP levels (Figure 6C).

**Figure 6:**
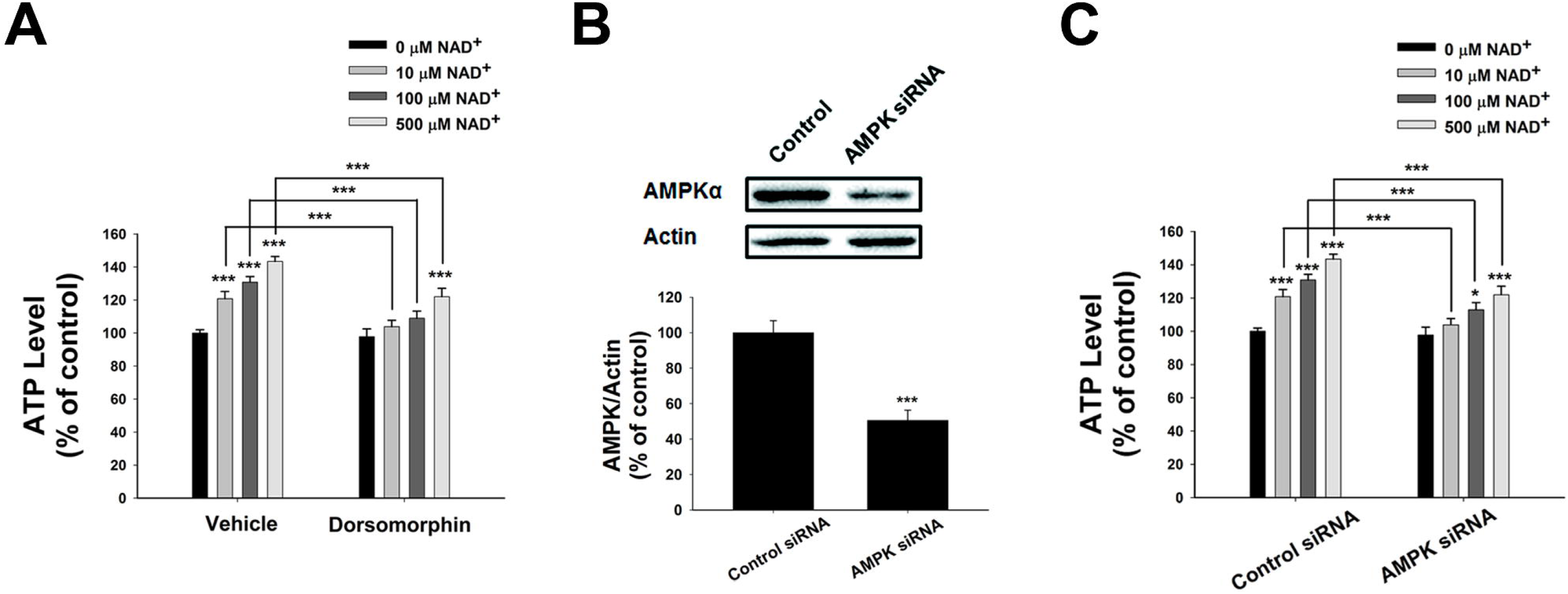
AMPK mediated the NAD^+^-induced increases in the intracellular ATP levels of BV2 microglia under basal conditions. (A) Dorsomorphin, an AMPK inhibitor, prevented NAD^+^-induced ATP increases in BV2 cells. The cells were co-treated with 15 μM dorsomorphin and NAD^+^ for 3 h before the ATP assay were conducted. (B) AMPK siRNA treatment led to a significant decrease in the AMPK protein level of BV2 cells. (C) AMPK siRNA significantly attenuated the NAD^+^-induced ATP increases of BV2 cells. The cells were pretreated with AMPK siRNA for 24 h. Subsequently the cells were treated with NAD^+^ for 3 h. N = 12. The data were pooled from four independent experiments. *, *P* < 0.05; **, *P* < 0.01; ***, *P* < 0.001.

### AMP treatment can increase both the intracellular adenylate levels and the AMPK activity of BV2 microglia under basal conditions

Our study has indicated the extracellular NAD^+^ can increase the intracellular AMP levels by enhancing the levels of intracellular adenosine that can be converted by adenosine kinase. We propose that the increased AMP may lead to increased levels of ATP and ADP as well as increased AMPK activity. To test the validity of this proposal, we determined the effects of AMP treatment, that has been reported to increase intracellular AMP levels (Brenner and Gorin, 1978), on the intracellular levels of ATP and ADP as well as the AMPK activity of BV2 microglia under basal conditions. Our study showed that AMP treatment led to a significant increase in the intracellular AMP levels of the cells (Figure 7A). The AMP treatment also produced significant increases in the intracellular levels of ATP and ADP (Figures 7B, 7C). Moreover, the AMP treatment significantly increased AMPK phosphorylation without affecting the intracellular AMP / ATP ratios of the cells (Figures 7D, 7E).

**Figure 7:**
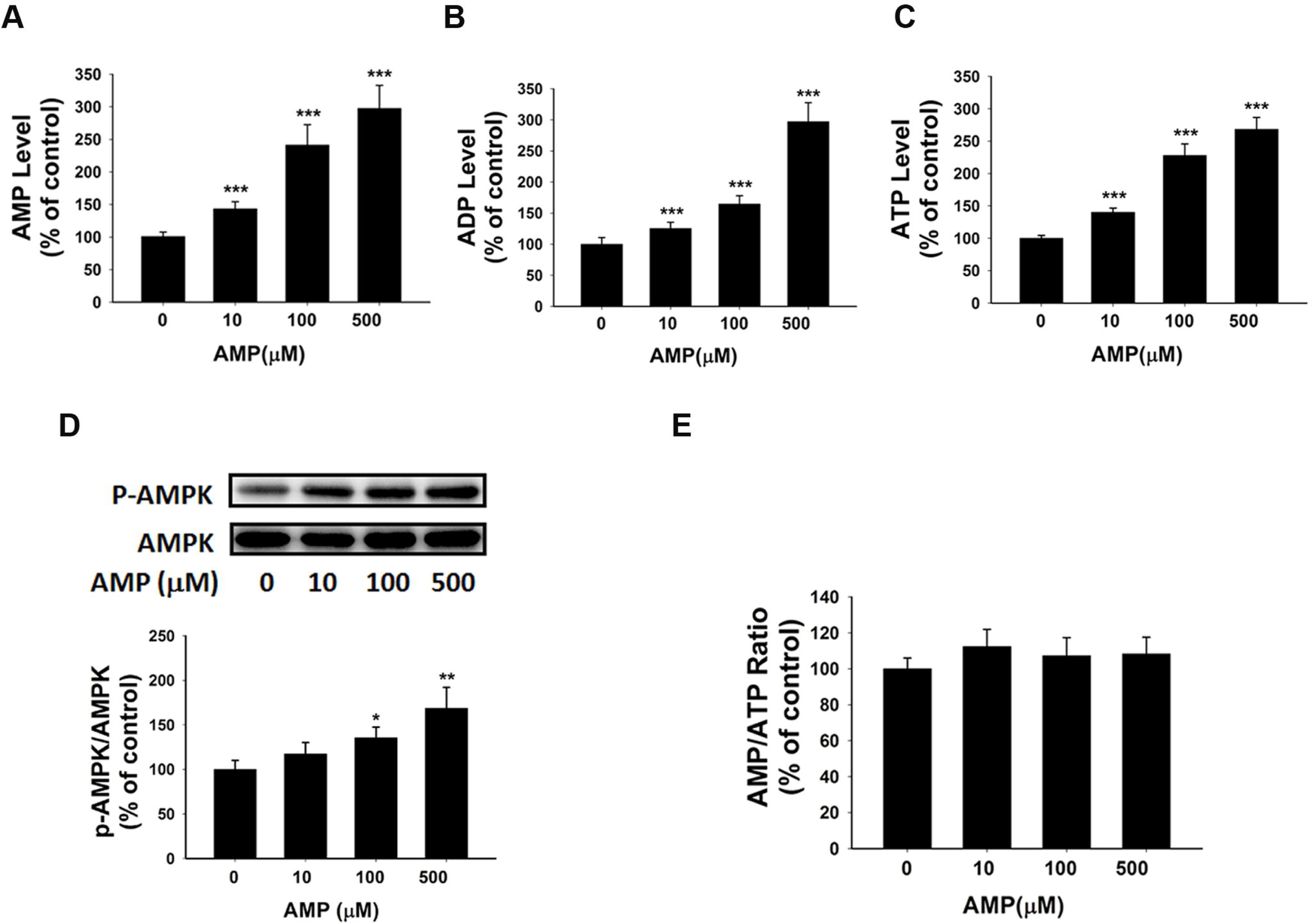
AMP treatment significantly increased the intracellular levels of ATP, ADP and AMP of BV2 microglia under basal conditions. (A) AMP treatment significantly increased the intracellular AMP level of the cells. (B) AMP treatment significantly increased the intracellular ADP level of the cells. (C) AMP treatment significantly increased the intracellular ATP level of the cells. (D) AMP treatment dose-dependently increased the AMPK phosphorylation of the cells. (E)AMP treatment did not influence the intracellular AMP / ATP ratios. The cells were treated with AMP for 3 h. Subsequently the assays were conducted. N = 16. The data were pooled from four independent experiments. *, *P* < 0.05; **, *P* < 0.01; ***, *P* < 0.001.

### Contribution of mitochondrial FoF1-ATP synthase to the NAD^+^-induced increases in the intracellular ATP levels of BV2 microglia under basal conditions

ADP is one of the major substrates of mitochondrial FoF1-ATP synthase, which is an important driving force for ATP synthesis (Stryer, 1995). Therefore, we proposed that the increased AMP-induced elevations of the intracellular ADP levels may enhance mitochondrial FoF1-ATP synthase activity, leading to increased intracellular ATP levels of BV2 microglia under basal conditions. We found that oilgomycin A, a selective inhibitor of mitochondrial FoF1-ATP synthase, completely blocked the NAD^+^-induced increases in the intracellular ATP levels (Figure 8).

**Figure 8:**
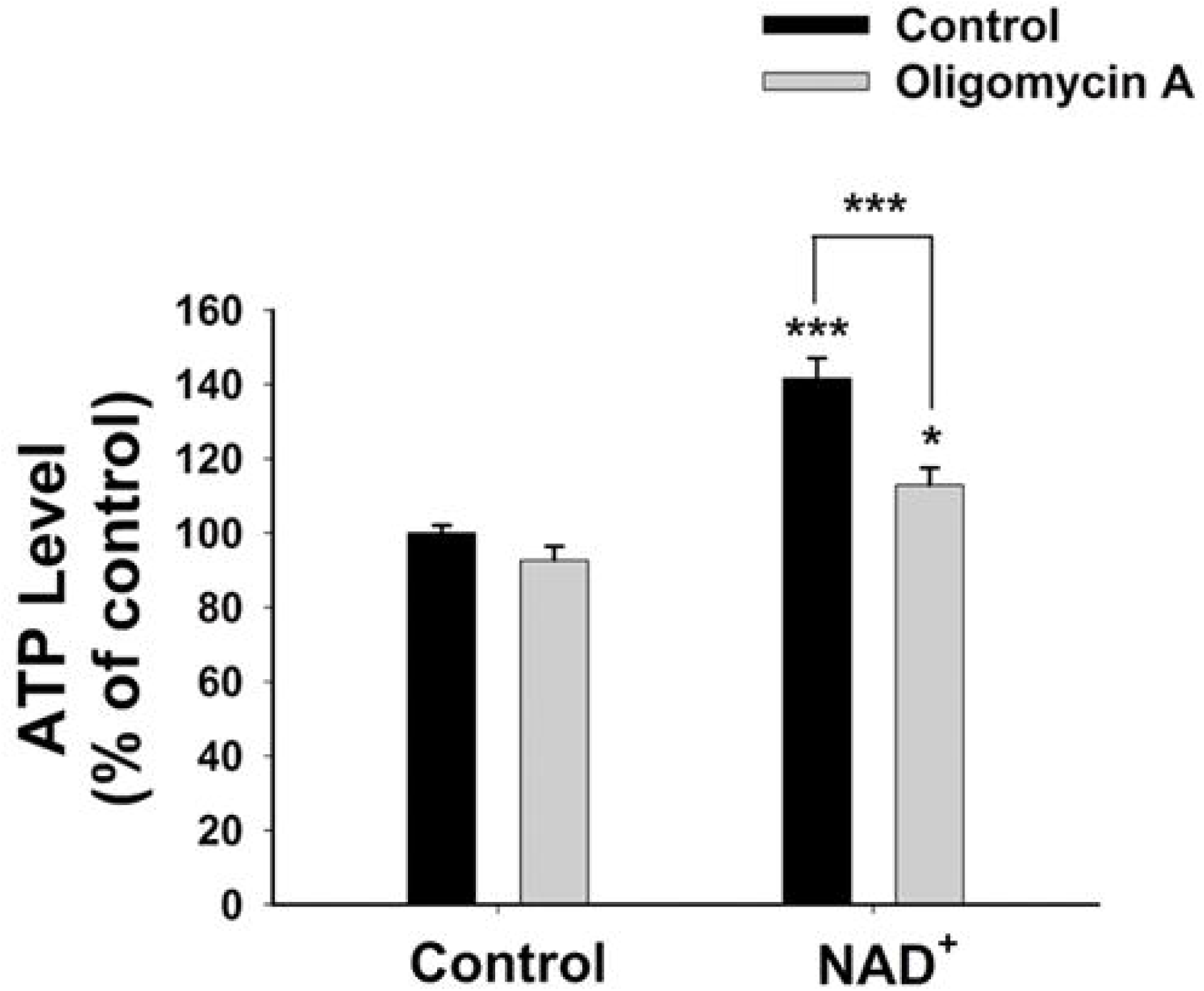
Mitochondrial FoF1-ATP synthase plays a significant role in the NAD^+^-induced increases in the intracellular ATP levels of BV2 microglia under basal conditions. Oligomycin A, a FoF1-ATP synthase inhibitor, prevented NAD^+^-induced ATP increases in BV2 cells. The cells were co-treated with 2 μM oligomycin A and 500 μM NAD^+^ for 3 h before the ATP assay were conducted.

## Discussion

The major findings of our current study includes: First, NAD^+^ treatment can significantly increase the intracellular levels of ATP, ADP and AMP of BV2 microglia under basal conditions. Second, our study has provided evidence arguing against the possibility that the NAD^+^ treatment increased the intracellular adenylate pool by affecting SIRT1, glycolytic rate or mitochondrial membrane potential of BV2 microglia under basal conditions. Third, blockage of ENTs can prevent the NAD^+^-induced increases in the intracellular adenylate pool of the cells. Fourth, NAD^+^ treatment can significantly increase the intracellular adenosine levels of the cells. Fifth, extracellular adenosine can produce increased intracellular adenylate levels of the cells. Sixth, adenosine kinase mediates the effects of NAD^+^ on the intracellular adenylate pool of the cells. Seventh, AMPK also mediates the NAD^+^-induced increases in the intracellular ATP. Eighth, inhibition of mitochondrial FoF1-ATP synthase can block the NAD^+^-induced increases in the intracellular ATP levels of the cells.

Previous studies have indicated that extracellular NAD^+^ significantly decreases the damage of the cells exposed to various pathological insults by such mechanisms as improving glycolysis, preventing mitochondrial depolarization and activating SIRT1 (Ying et al., 2003; Alano et al., 2004; Pillai et al., 2005; Ying, 2008; Alano et al., 2010; Zhang and Ying, 2018). However, our study has shown that the treatment of BV2 microglia with 0.01 – 0.5 mM NAD^+^ did not significantly increase the glycolytic rate and mitochondrial membrane potential of BV2 microglia under basal conditions. Moreover, our study did not show that SIRT1 inhibition can affect the NAD^+^-produced increases in the intracellular adenylate pool of the cells under basal condition. Therefore, our study has indicated that the mechanisms underlying the NAD^+^ treatment-produced effects on the cells under pathological insults are not applicable to the cells under basal conditions. These observations are not surprising for the following reasons: The cytosolic NAD^+^ concentrations are normally in the ranges between 1 – 10 mM (Ying, 2008). Because NAD^+^ enters cells by gradient-driven transport (Alano et al., 2010), the NAD^+^ at the concentrations between 0.01 – 0.5 mM, which was used in our study, should not be able to enter the cells to influence the adenylate pools of BV2 microglia under basal conditions.

Our current study has indicated that extracellular NAD^+^ produces its biological effects on cells under basal conditions through mechanisms that are distinctly different from the mechanisms found in the cells exposed to various pathological insults. Our study has indicated that extracellular NAD^+^ can produce significant biological effects on BV2 microglia under basal conditions through the following mechanism (Fig. 9): Extracellular NAD^+^ is degraded into adenosine that enters the cells through ENTs, which is converted to AMP by adenosine kinase. Increased AMP can lead to both increased AMPK activity and increased intracellular ADP levels - a major driving force for mitochondrial FoF1-ATP synthase, which jointly produce the increased intracellular ATP levels of BV2 cells under basal conditions.

**Figure 9:**
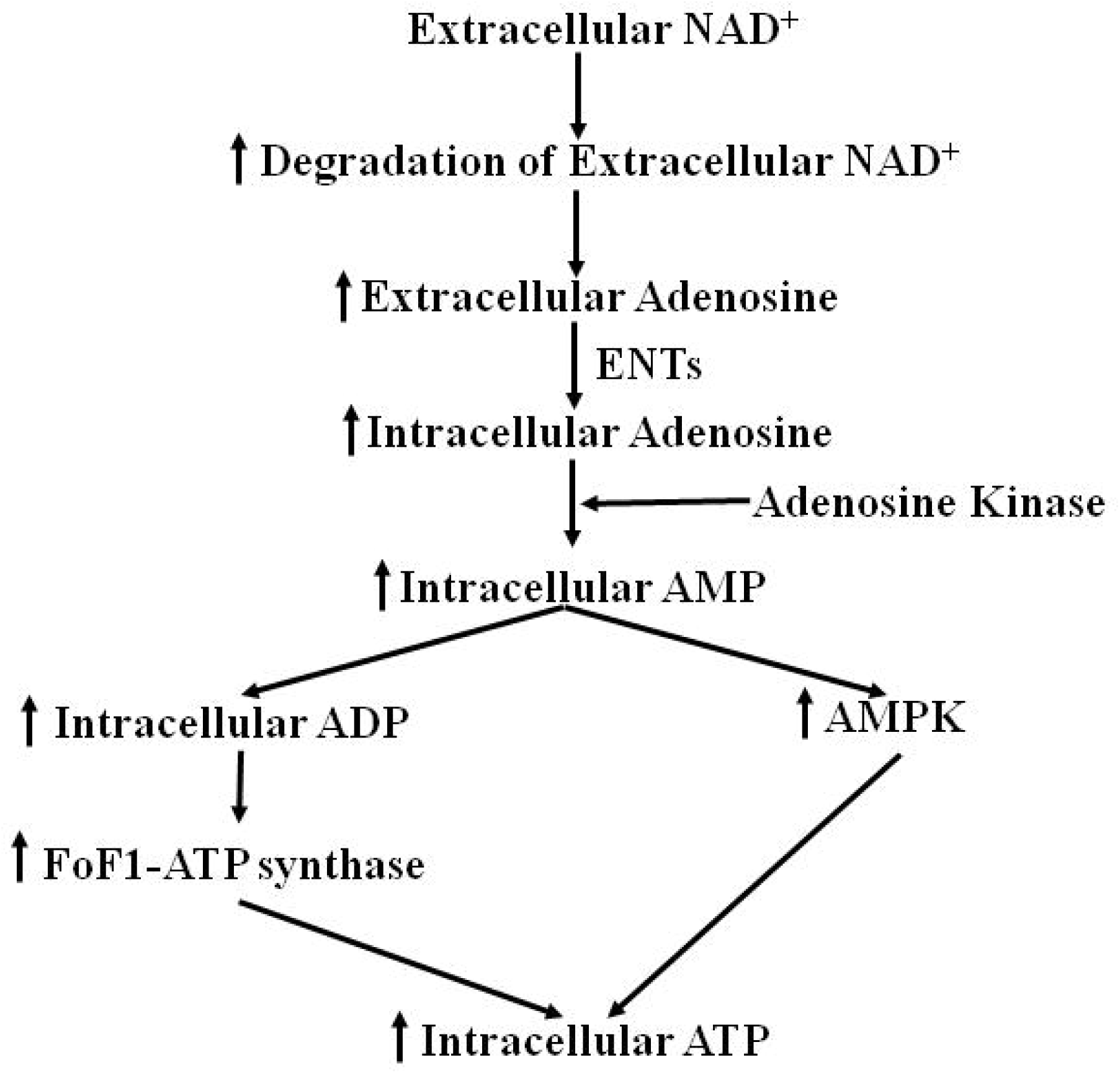
Diagrammatic presentation of the mechanisms underlying the effects of NAD^+^ on the intracellular adenylate pool of BV2 microglia under basal conditions.

Although the NAD^+^ at the concentrations between 0.01 – 0.5 mM can not directly enter cells to increase intracellular NAD^+^ levels, we still found that the NAD^+^ can significantly increases the intracellular levels of NAD^+^ and adenylate of BV2 microglia through extracellular degradation into adenosine. Since intracellular adenosine levels are normally in the nanomolar range (Latini and Pedata, 2001), the extracellular NAD^+^-generated adenosine can enter cells to increase intracellular adenosine levels thus increasing the intracellular adenylate levels through the activities of adenosine kinase and AMPK. In other words, the exceedingly low intracellular adenosine levels under normal physiological conditions is an unique ‘attractor’ and base for the relatively low concentrations of extracellular NAD^+^ to produce its significant biological effects on cells. While a previous study reported that extracellular NAD^+^ can be degraded into adenosine, which is transported into astrocytes through ENTs (Latini and Pedata, 2001), to our knowledge, our current study has provided the first direct evidence showing that extracellular adenosine generated from extracellular NAD^+^ degradation mediates the biological effects of exogenous NAD^+^ by adenosine kinase- and AMPK-mediated pathways.

Our study has shown that relatively low concentrations of NAD^+^ can increase both extracellular and intracellular adenosine levels, the AMPK activity and intracellular adenylate pools, all of these factors have been shown to enhance defensive potential of both normal cells and stressed cells against toxic insults: AMPK activation has been reported to produce multiple beneficial effects for both normal and stressed cells and tissues, for examples, AMPK activation can expand the lifespan in multiple model organisms (Apfeld et al., 2004; Stenesen et al., 2013); and AICAR, an AMPK activator, protects the liver from fatty changes associated with chronic alcohol use in rats (Tomita et al., 2005); adenosine can produce beneficial effects under multiple pathological conditions, including myocardial ischemia, seizure and acute kidney injury (Ely and Berne, 1992; Zhang et al., 1993; Yap and Lee, 2012); and an increase in the adenylate pool and absolute ATP concentrations could enhance the potential of cells to fulfill useful work (Ataullakhanov and Vitvitsky, 2002), and an intracellular ATP level is a switch for the decision between apoptosis and necrosis (Leist et al., 1997). Collectively, our current study has suggested a novel mechanism to account for the profound protective effects of NAD^+^ administration in the animal models of a number of diseases and aging (Liu et al., 2014; Scheibye-Knudsen et al., 2014; Fang et al., 2016): In addition to its reported capacity to enhance the capacity of the cells under pathological insults to defend against the cell death-inducing insults, the extracellular NAD^+^ may also increase the defensive potential of the normal cells that have not been attacked yet at the time of the exposures to exogenous NAD^+^ by increasing both extracellular and intracellular adenosine levels, the activities of adenosine kinase and AMPK, and intracellular adenylate pools.

It is noteworthy that the intravenous NAD^+^ administration in all of the animal studies should lead to the NAD^+^ concentrations that are well below milimolar range (Ying et al., 2007; Wang et al., 2014; Zhang et al., 2016; Xie et al., 2017). Since the cytosolic concentration of NAD^+^ is normally in the range between 1 – 10 mM, it is usually assumed that the NAD^+^ administration-produced NAD^+^ concentrations in the blood should not be able to enter cells to produce biological effects. However, our study has shown that as low as 10 μM NAD^+^ can lead to significant increases in the ATP levels of all of the cell types we have studied on this topic, including BV2 microglia, PC12 cells and C6 glioma cells (Zhang et al., 2018). Therefore, our findings have general value for understanding the mechanisms underlying the biological effects of NAD^+^ administration in models of diseases, aging or healthy controls: The relatively low concentrations of extracellular NAD^+^ can still produce its profound effects on cells through its extracellular degradation into adenosine, which may lead to increased intracellular levels of adenosine, AMP, ADP and ATP on the basis of the activities of ENTs, adenosine kinase, AMPK and mitochondrial FoF1-ATP synthase.

ATP plays critical roles in microglia: Basal ATP release is a critical biological signal of microglia; and extracellular ATP induces migration, chemotaxis and ramification of microglia (Honda et al., 2001; Wollmer et al., 2001; Duan et al., 2009; Ma et al., 2014). Therefore, our findings regarding the capacity of NAD^+^ treatment to enhance the intracellular ATP levels of BV2 microglia under basal conditions also have the following implications: NAD^+^ administration could produce significant biological impact on microglial activities under both pathological conditions and normal physiological conditions.

## Acknowledgment

This study was supported by major basic research grants from Science and Technology Commission of Shanghai Municipality, Grant No.16JC1400500 and No.16JC1400502 (to W.Y.), and Chinese National Science Foundation Grants #81271305(to W.Y.).

## Legends of Supplemental Figures

**Supplementary Figure 1. NAD^+^ treatment did not significantly affect the glycolytic rate or mitochondrial membrane potential of BV2 microglia under basal conditions.** (A) 2-NBDG uptake assay did not show that the NAD^+^ treatment significantly affected the glycolytic rate of BV2 cells. (B) FACS-based JC-1 assay did not show that the NAD^+^ treatment affected the mitochondrial membrane potential of the cells. (C) Quantifications of the results from the FACS-based JC-1 assay did not show that the NAD^+^ treatment significantly affected the mitochondrial membrane potential of the cells. The cells were treated with NAD^+^ for 3 h. Subsequently the assays were conducted. (D) The Complex I inhibitor rotenone was not capable of preventing 500 μM NAD^+^-induced increases in the intracellular ATP levels. (E) The Complex IV inhibitor sodium azide was not capable of preventing 500 μM NAD^+^-induced increases in the intracellular ATP levels. N = 12. The data were pooled from four independent experiments.

**Supplementary Figure 2.** Ten μM EX527, a selective SIRT1 inhibitor, did not affect the AMPK phosphorylation induced by 500 μM NAD^+^ of BV2 microglia under basal condition. The cells were co-treated with NAD^+^ and EX527 for 3 h. Subsequently Western blot assays were conducted. N = 12. The data were pooled from three independent experiments. *, *P* < 0.05; **, *P* < 0.01; ***, *P* < 0.001.

**Supplementary Figure 3. Nicotinamide treatment did not affect the intracellular ATP levels of BV2 microglia under basal condition.** Treatment of the cells with 10 or 100 μM nicotinamide did not affect the intracellular ATP levels, while treatment of the cells with 500 μM nicotinamide slightly increased the intracellular ATP levels. The cells were treated with nicotinamide for 3 h. Subsequently ATP assays were conducted. N = 12. The data were pooled from three independent experiments. *, *P* < 0.05.

